# Consistent Neural Representation of Affective States between Watching and Recall using fNIRS

**DOI:** 10.1101/2025.11.22.689961

**Authors:** Chaery Park, Jongwan Kim

## Abstract

People tend to recall similar feelings when remembering the same event. Yet the neural mechanisms by which emotional valence and arousal are represented during perception and later recall remain unclear. Here, we used functional near-infrared spectroscopy to record prefrontal cortex activity while participants viewed and then freely recalled short video clips designed to elicit four affective states (amusement, calmness, disgust, and sadness). Multidimensional scaling revealed that both modality-general (shared across viewing and recall) and modality-specific (unique to each phase) dimensions of valence and arousal were significant, and cross-modal classification successfully decoded valence and arousal. Our results support a hierarchical model in which a stable core affect is preserved across perception and memory, while context-dependent refinements adapt emotional representations during recall. This work highlights fNIRS as a useful tool for studying emotional memory in ecologically valid settings and lays groundwork for multimodal imaging of affective replay.

## 1. Introduction

Emotion plays a central role in both perception and memory, especially when the stimuli carry high emotional salience. Emotionally charged stimuli are typically encoded more vividly and remembered more accurately (Kensinger & Schacter, 2008). Yet, it remains unclear whether the emotional response elicited during recall mirrors the emotion originally experienced during the perception of the stimulus. While emotionally salient events evoke robust physiological and neural responses at the time of perception, those responses may be altered or reconstructed during recall depending on context and internal states (Pascalis et al., 2013). Understanding the degree of neural similarity between emotional responses during perception and during recall represents a key challenge in emotion and memory research. The present study addresses this gap by directly comparing the neural representations of emotional experiences during viewing and recall of emotionally evocative video stimuli. The goal is not merely to examine whether participants remember the same content, but to assess whether the emotional tone of the original experience is preserved or transformed during recall.

Chen et al. (2017) demonstrated using fMRI that participants who viewed and then freely recalled the same video stimuli exhibited highly similar scene-specific activation patterns in high-level associative networks, including the default mode network. Importantly, these authors argued that perceived content is not simply replayed during recall, but is systematically transformed and reconstructed in memory. This suggests that emotional experiences may be reconstructed in memory in ways that are both personal and socially shared. High-level emotional and cognitive processes are known to involve areas such as the medial prefrontal cortex and the posterior parietal cortex (Lindquist et al., 2012), and these regions may also play a central role during the recall of emotional memories.

Previous studies have predominantly relied on fMRI, which offers high spatial resolution but has limitations when it comes to capturing naturalistic emotional responses and verbal recall. Participants in fMRI studies must remain still in a confined space, view stimuli within a narrow field of view, and often hear auditory stimuli through headphones under noisy conditions. These physical constraints reduce ecological validity and may hinder the measurement of natural emotional expression and free recall. In contrast, functional near-infrared spectroscopy (fNIRS) provides a promising alternative that addresses many of these limitations. fNIRS is relatively tolerant to motion artifacts, allowing participants to speak and gesture naturally during measurement. Its open design enables testing in realistic environments, and the system is more affordable and portable than fMRI. It is also quiet and allows for real-time monitoring of brain activity, making it well suited for emotionally engaging tasks conducted in ecologically valid settings (Quaresima & Ferrari, 2019).

This study is grounded in converging neuroscientific evidence that the cognitive evaluation and regulation of emotion are implemented in the prefrontal cortex—specifically dorsolateral, ventrolateral, and ventromedial/orbitofrontal subdivisions (dlPFC, vlPFC, vmPFC/OFC). Prefrontal regions allocate attention to affective cues, assign value to those cues (valuation), and implement reappraisal strategies, thereby modulating both peripheral physiological responses and the subjective intensity of emotional experience (Berboth & Morawetz., 2021; Denny et al., 2023). In addition, functional lateralization in the frontal cortex—where left frontal activity is more strongly associated with approach motivation and positive affect, and right frontal activity with withdrawal motivation and negative/aversive processing—supports the expectation that physiological markers distinguishing positive versus negative emotion will be particularly salient over prefrontal sites (Harmon-Jones & Gable, 2018). Building on these considerations, and on mechanistic evidence that dlPFC and vlPFC make dissociable contributions to reappraisal versus distraction and related control operations (Mo et al., 2023), and that vmPFC encodes value signals that are sensitive to affective context (Chakravarthula & Padmala., 2023), we preregistered the prefrontal montage as our primary region of interest (ROI). Accordingly, we specified two a priori hypotheses: (1) prefrontal activity would differ between positive and negative emotion conditions; and (2) lateralization metrics (left–right asymmetry) would show condition effects, with all measurements acquired bilaterally and analyzed with hemisphere as an explicit factor (Berboth & Morawetz., 2021; Harmon-Jones & Gable, 2018; Denny et al., 2023).

Given these characteristics, the present study used fNIRS to examine differences in prefrontal activation during the viewing and recall of emotional video stimuli under ecologically valid conditions. Specifically, we investigated how closely emotional experiences during perception resemble those during recall at the neural level. Furthermore, by assessing whether multiple participants show similar prefrontal activation patterns during the recall of the same stimuli, we aimed to provide insight into the shared nature of emotional memory. We expect that our fNIRS-based approach will contribute to a more realistic and detailed understanding of emotional memory as it occurs in everyday contexts.

## 2. Methods

### 2.1. Participants

A total of 36 participants (7 male, 29 female) took part in the study. Due to data acquisition issues, data from 4 participants were excluded. The final analysis included 32 participants (6 male, 26 female), aged 19 to 26 years. The study protocol was approved by the Institutional Review Board of the affiliated university (Approval No. 2025-02-009-001).

### 2.2. Stimuli and apparatus

Prior to the main experiment, a pilot study was conducted with seven participants to select appropriate video stimuli. A total of 38 videos were collected from YouTube, and participants rated each video on a 7-point scale across 15 emotion adjectives: sad, anxious, gloomy, angry, irritated, fearful, disgusted, surprised, relaxed, calm, peaceful, satisfied, cheerful, joyful, and happy. For each emotional condition, we averaged the ratings of the relevant adjectives and selected the three videos with the highest mean scores as the stimuli for the main experiment (e.g., for the negative/low-arousal condition, the average of “sad,” “anxious,” and “gloomy” ratings was used for selection).

In the main experiment, participants were presented with three short videos (ranging from 29 to 79 seconds) for each condition (Table 1). To ensure accurate measurement in naturalistic conditions, we used the NIRSIT LITE system (OBELAB Inc.), a portable and validated device for fNIRS-based research (Baker et al., 2017).

**Table 1.**
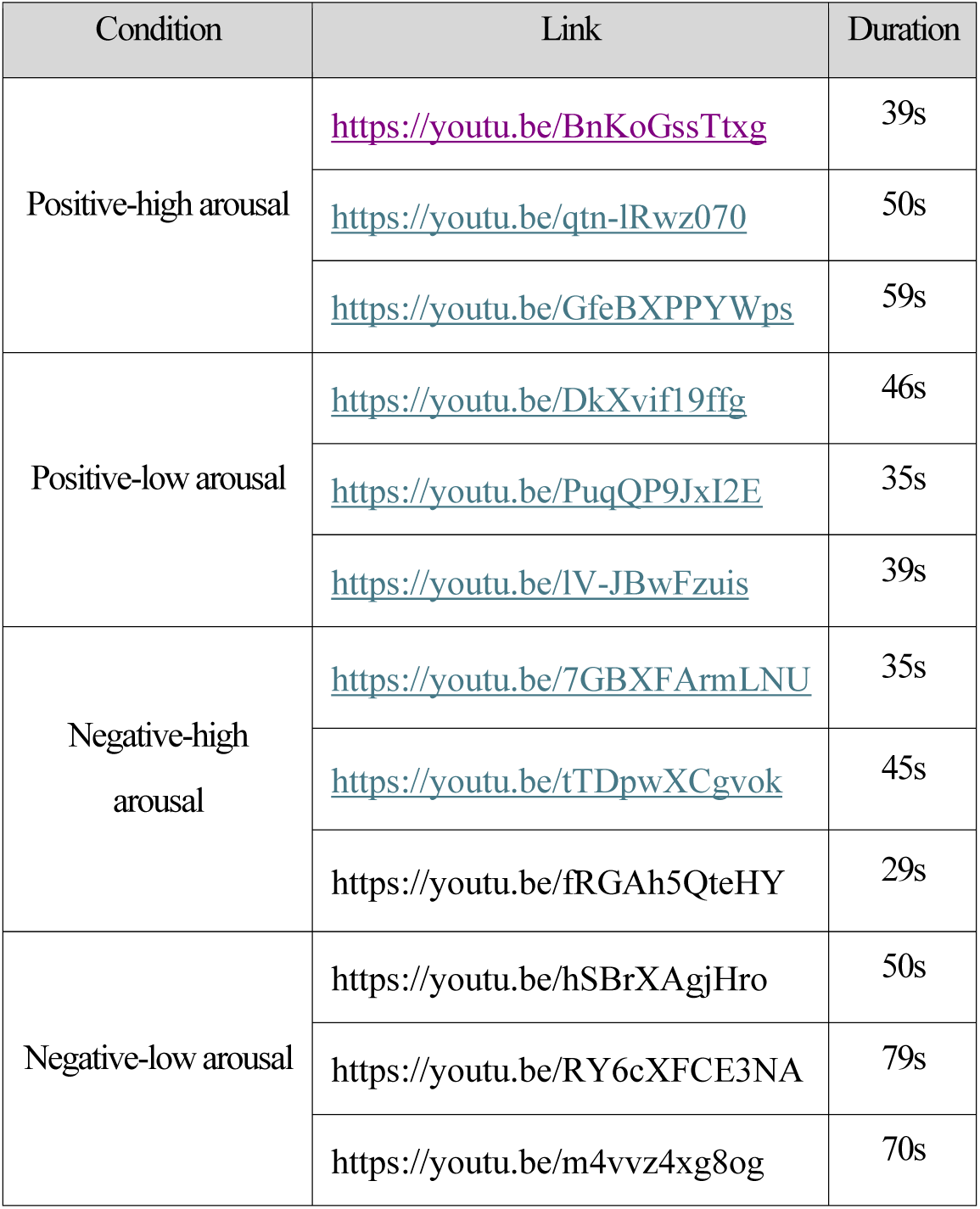
Stimuli selected for main experiment.

**Table 2.** Estimated AAL coordinates and region labels.

| Channel | Hemisphere | x | y | z | AAL index | Region Label |
| --- | --- | --- | --- | --- | --- | --- |
| 1 | Right | 54.67 | 44 | 0 | 10,12 | Inferior frontal Gyrus |
| 2 | Right | 48 | 52.67 | 12.67 | 6, 10 | Middle Frontal Gyrus |
| 3 | Right | 36.33 | 58.33 | 24.67 | 6 | Middle Frontal Gyrus |
| 4 | Right | 39 | 64.67 | 0 | 6 | Middle Frontal Gyrus |
| 5 | Right | 26.33 | 69.67 | 12.67 | 4 | Superior Frontal Gyrus,<br>Dorsolateral |
| 6 | Right | 26.33 | 69.67 | 24.67 | 4, 20 | Superior Frontal Gyrus, Medial |
| 7 | Right | 14.33 | 73 | 0.33 | 4, 20, 22 | Superior Frontal Gyrus, Medial |
| 8 | Center | 0.33 | 68 | 11.33 | 19, 20 | Superior Frontal Gyrus, Medial |
| 9 | Left | -13 | 68 | 24 | 3, 19 | Superior Frontal Gyrus, Medial |
| 10 | Left | -14.33 | 73 | 0 | 3, 19, 21 | Superior Frontal Gyrus, Medial |
| 11 | Left | -26 | 68 | 12 | 3 | Superior Frontal Gyrus,<br>Dorsolateral |
| 12 | Left | -35.67 | 57.67 | 23.67 | 5 | Middle Frontal Gyrus |
| 13 | Left | -38 | 63 | 0 | 5 | Middle Frontal Gyrus |
| 14 | Left | -45.67 | 51.67 | 12.33 | 5, 9 | Middle Frontal Gyrus |
| 15 | Left | -52.33 | 43.67 | -0.33 | 9, 11 | Inferior Frontal Gyrus |

The NIRSIT LITE device contains 15 channels focused on the prefrontal cortex. The raw fNIRS signals were converted into oxyhemoglobin concentrations using the Modified Beer–Lambert Law (MBLL), and low-frequency physiological noise (e.g., respiration and heartbeat) was attenuated using a low-pass filter. Motion artifacts were corrected using an onboard motion sensor.

Each channel was mapped to specific subregions of the prefrontal cortex, estimated using the Montreal Neurological Institute (MNI) coordinates and the AAL atlas (AAL3v1; Rolls et al., 2020). These regions are functionally linked to high-level cognitive and emotional processes. For instance, the inferior frontal gyrus (IFG) is associated with inner speech and emotional inhibition (Aron et al., 2004), the middle frontal gyrus (MFG) with working memory and attentional switching (D’Esposito et al., 1995), the dorsolateral superior frontal gyrus (dlPFC) with self-regulatory emotional control (Ochsner et al., 2004), and the medial superior frontal gyrus (mPFC) with autobiographical memory and emotional introspection (Buckner & Carroll, 2007). This layout enabled comprehensive measurement of prefrontal responses to emotional stimuli during both induction and recall phases.

### 2.3. Procedure

Prior to the experiment, participants were carefully screened to ensure that their physiological states such as heart rate and respiration were not influenced by external factors. They were instructed to refrain from alcohol consumption on the day of the experiment and to avoid wearing excessive makeup on their foreheads, as the sensors needed to be directly attached to the skin. Upon arrival at the laboratory, participants received a briefing and provided written informed consent.

To ensure the accuracy of light detection, the ambient lighting was adjusted to minimize external interference. The experiment was conducted in a controlled setting to block environmental light. The fNIRS sensors were carefully attached to the forehead, and their placement was repeatedly checked. Hair in the measurement area was parted, and the skin was cleaned with medical-grade alcohol swabs if necessary, ensuring secure contact between the sensor and the skin.

A block design, commonly used in fNIRS studies, was employed. This design allows for simple experimental structure and repeated stimulus presentation, enhancing the reliability of signal measurement. The experiment consisted of two sessions: a watching session and a recall session. In the watching session, participants viewed 12 short emotional video clips, with three clips for each of the four emotional conditions. The order of the conditions was pseudorandomized, and the order of the three trials within each condition was also randomized to reduce potential order effects and stimulus predictability. After the watching session, a short break of approximately 3 minutes was provided to relieve any discomfort from sensor placement.

In the recall session, participants recalled the same 12 clips in the identical order as in the watching session. At the start of each trial, a still image from the first frame of the original video was shown to cue recall. Participants were then asked to verbally describe the scene freely for one minute. Although participants were allowed to move their bodies and heads freely to promote natural speech, they were instructed to minimize movement as much as possible to preserve signal quality. The recall session consisted of 12 trials in total, and the full experiment lasted approximately 50 minutes. A brief debriefing followed upon completion of the study.

### 2.4. Preprocessing

The raw fNIRS data collected in this study were preprocessed using OBELAB NIRSIT-QUEST software. First, channels with five or fewer consecutive invalid values were corrected using nearest-neighbor interpolation. If more than five consecutive invalid values were detected, the entire channel was excluded from further analysis. Channels with a median intensity value below 30 were also excluded, as such low-intensity signals are considered unreliable (Yücel et al., 2021). Channels were further removed if the coefficient of variation exceeded 15%, indicating unstable signal quality (Pfeifer et al., 2018), or if more than 5% of the entire time series consisted of identical values, suggesting signal saturation.

Remaining data were converted from light intensity to optical density (OD) (Yücel et al., 2021). The OD signals were then corrected for motion artifacts using the Temporal Derivative Distribution Repair (TDDR) method (Fishburn et al., 2019). Finally, the optical density data were converted to concentrations of oxy- and deoxy-hemoglobin using the Modified Beer–Lambert Law (MBLL), based on the methods of Delpy et al. (1988) and Cope & Delpy (1988). In accordance with ISO guidelines, differential pathlength factors were not applied. Hemoglobin concentrations were expressed in units of mm·mM, and molar extinction coefficients were adopted from the optimized dataset reported by Zhao et al. (2017).

In the final preprocessing step, missing channels from the exclusion steps were reconstructed using rejection padding—where the signal was replaced with the mean value of neighboring channels. This preprocessing pipeline, grounded in methodological recommendations from prior fNIRS studies (Scholkmann et al., 2014), ensured that the resulting prefrontal signal patterns reflected robust and biologically meaningful hemodynamic activity.

### 2.5. Model specification

Following preprocessing, a GLM was specified in line with best-practice recommendations (Yücel et al., 2021). Beta coefficients were estimated using the precoloring method to address temporal autocorrelations, and Wavelet-MDL was employed to remove slow trends. Hemodynamic response modeling incorporated a canonical HRF kernel for temporal smoothing (Tak et al., 2008).

Condition-specific regressors were constructed for each emotional category. Temporal and dispersion derivatives of each regressor were included to accommodate variability in onset and duration of the hemodynamic responses. To further account for potential noise, nuisance regressors were added. These included motion estimates obtained from the onboard IMU sensor, capturing head movement angles in the x, y, and z axes (Penny et al., 2011). In addition, short-separation signals from optode pairs with source-detector distances ≤8 mm were averaged and regressed out of each channel’s time series to mitigate the effects of superficial systemic noise (Gagnon et al., 2011).

The specified model was fit separately to oxyhemoglobin (HbO), deoxyhemoglobin (HbR), and total hemoglobin (HbT) time series in each channel. For consistency in interpretation, regressors fit to HbR were mirror-reversed so that positive beta coefficients consistently indicated activation across all hemoglobin types. Contrast-based beta estimates were computed on a per-channel basis, and region-of-interest (ROI) level summaries were obtained by averaging these beta values across anatomically grouped channels. In addition to the GLM, a finite impulse response (FIR) model was used to estimate time-course dynamics over a 24-second window post-stimulus onset. This was achieved using 24 epoch regressors (one per second), allowing fine-grained temporal tracking of the hemodynamic response (Henson, 2002; Brett et al., 2002).

For the following statistical analyses, a regressor for each of the 24 trials was entered into the GLM. Patterns of activation were formed by beta estimates for each of 24 trials scaled by the square root of the estimated residual variance for each channel (Misaki et al., 2010). Furthermore, the data for each trial were standardized across channels to have zero mean and unit variance (Pereira et al., 2009).

### 2.6. Statistical analyses

#### 2.6.1. Multidimensional scaling (MDS)

Multidimensional scaling (MDS) is a statistical technique used to represent complex patterns of similarity or dissimilarity among stimuli or participants within a reduced-dimensional space. This technique enables the visualization of latent structural relationships embedded in the data. In the current study, MDS was applied to examine how the 24 emotional trials across both watching and recall sessions are organized in a low-dimensional space, with the expectation that the resulting dimensions would reflect core affective components such as valence and arousal.

Since MDS does not inherently provide statistical inference, the resulting coordinates were subjected to Procrustes rotation (Gower, 1975) to align the MDS solution with a predefined design matrix. The correspondence between the rotated coordinates and the design was then assessed by calculating the correlation between them.

To perform MDS, a 24 × 24 stimuli-by-stimuli correlation matrix was computed for each participant based on their fNIRS activation patterns. These individual matrices were then averaged to yield a group-level similarity matrix, which was subjected to five-dimensional classical MDS. The resulting spatial configuration was rotated via Procrustes transformation to maximize its alignment with theoretical design contrasts.

The design matrix was constructed to capture both modality-general and modality-specific representations of affective experience (Table 3) (Shinkareva et al., 2016; Kim et al., 2017). This approach allows us to determine whether emotion is encoded similarly across sensory modalities or in a modality-dependent manner. For modality-general valence, we applied a contrast vector where positive and negative stimuli across both modalities were coded as [1, –1, 1, –1], reflecting a shared representation of affect regardless of sensory modality. In contrast, modality-specific valence was derived by multiplying the sensory-modality contrast [1, 1, –1, –1] with the shared valence contrast, yielding [1, –1, –1, 1]. This pattern represents a scenario in which the two modalities encode emotional valence in a mirrored or opposing fashion, thereby supporting the notion of modality-specific representation.

**Table 3.** Design matrix of experimental conditions.

| Modality | Valence | Arousal | Dimension 1: Modality | Dimension 2: Modality-general valence | Dimension 3: Modality-general arousal | Dimension 4: Modality-specific valence | Dimension 5: Modality-specific arousal |
| --- | --- | --- | --- | --- | --- | --- | --- |
| Watching | Positive | High | 1 | 1 | 1 | 1 | 1 |
|  |  | Low | 1 | 1 | -1 | 1 | -1 |
|  | Negative | High | 1 | -1 | 1 | -1 | 1 |
|  |  | Low | 1 | -1 | -1 | -1 | -1 |
| Recall | Positive | High | -1 | 1 | 1 | -1 | -1 |
|  |  | Low | -1 | 1 | -1 | -1 | 1 |
|  | Negative | High | -1 | -1 | 1 | 1 | -1 |
|  |  | Low | -1 | -1 | -1 | 1 | 1 |

From this analysis, we hypothesize that if valence is represented in a modality-specific manner, the spatial distribution of stimuli from one modality would resemble a flipped version of the distribution from the other modality within the MDS space. This contrast-based framework provides a formal test of whether emotional representations are shared across modalities or uniquely structured within each sensory channel.

#### 2.6.2. Classifications

To evaluate whether affective states including valence, arousal (2-way), and emotion category (4-way) could be predicted based on fNIRS activation patterns, classification analyses were conducted using support vector machine (SVM) classifiers. Two types of classification were performed: cross-participant within-modal classification and cross-participant cross-modal classification.

In the within-modal classification, data from one participant were held out as the test set, while the remaining participants’ data from the same session (either watching or recall) were used to train the classifier. This leave-one-subject-out cross-validation was repeated until each participant had been used once as the test subject. The average classification accuracy across folds was computed for each affective dimension. This procedure was conducted separately for the watching and recall sessions. Above-chance classification performance would suggest that affective neural responses are consistent across individuals, indicating that emotional state representations can be generalized from one participant to another within the same modality (i.e., within either watching or recall). This consistency implies shared neural encoding of emotional experiences that is stable across individuals when exposed to the same stimuli.

In the cross-modal classification, data from one participant in one session (e.g., watching) were used as the test set, while the remaining participants’ data from the other session (e.g., recall) were used to train the classifier. This process was repeated for each participant and for both directions, watching to recall and recall to watching. Successful classification in this cross-modal context would indicate that emotional states are represented similarly across modalities, implying that the neural encoding of an emotional state during passive viewing resembles the encoding during active recall.

Specifically, consistent patterns across modalities suggest that certain brain activation signatures for valence or arousal are modality-invariant and can be generalized across perceptual and memory-driven conditions. Such findings would support the idea of a shared neural representational space for emotional experience, independent of whether it is externally triggered or internally reconstructed.

Together, the classification analyses test whether affective responses can be robustly decoded across individuals and across task contexts. Significant classification accuracy in within-modal analyses supports individual-invariant encoding of emotional states, while above-chance performance in cross-modal analyses provides evidence for modality-general neural representations of affect.

These findings contribute to our understanding of how emotional experiences are neurally encoded and generalized across people and across perceptual versus memory-based contexts.

## 3. Results

### 3.1. MDS

Affective video stimuli from both the watching and recall sessions were represented within a five-dimensional solution derived from MDS (Fig. 1). Pearson correlation analyses were conducted to examine the correspondence between each dimension of the Procrustes-rotated MDS solution and the theoretical design matrix.

**Fig. 1.**
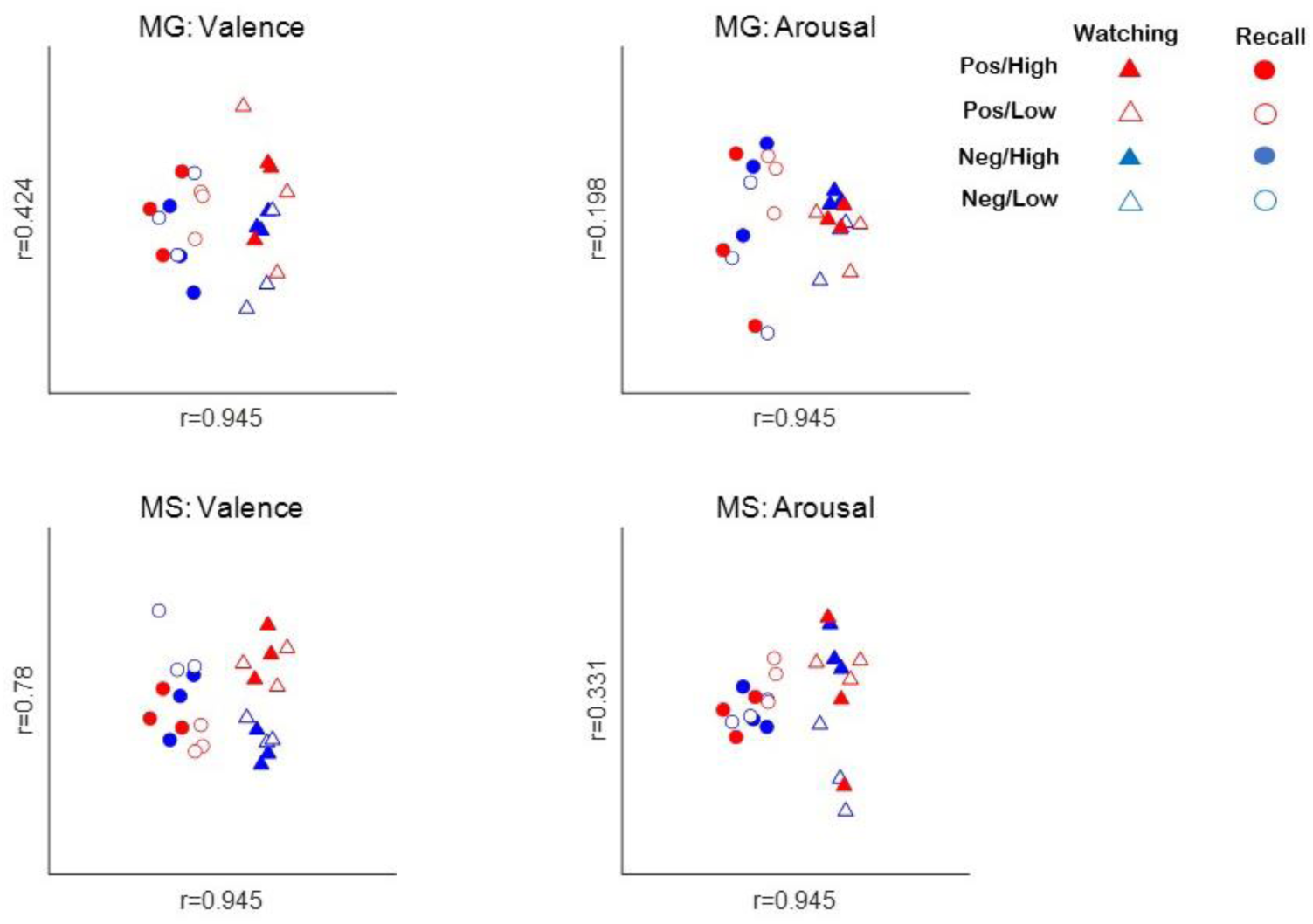
The results of rotated 5-dimensional MDS of the affective videos. r = Pearson correlation coefficient between the coordinates of the rotated MDS solution and the design values. MG: modality-general, MS: Modality-specific, Pos/High= Positive High, Pos/Low=Positive Low, Neg/High=Negative High, Neg/Low=Negative Low.

The first dimension, corresponding to modality, was significant (r = .945, p < .001), indicating substantial differences in fNIRS activation patterns between the watching and recall sessions. The second dimension, reflecting modality-general valence, was also significant (r = .424, p = .039), suggesting a consistent distinction between positive and negative videos across both sessions. The third dimension, representing modality-general arousal was not significant (r = .198, p = .354), indicating unclear separation between high- and low-arousal stimuli independent of modality. The fourth dimension, associated with modality-specific valence, was significant (r = .78, p < .001), suggesting that the representation of valence differed systematically between watching and recall. Finally, the fifth dimension, reflecting modality-specific arousal, was not significant (r = .331, p = .114), indicating that arousal-related representations were not modality general nor modality specific.

These results suggest that while core affective valence is consistently encoded across perceptual and mnemonic experiences, there also exist modality-specific representations, indicating partial transformation of emotional information between watching and recall.

### 3.2. Classifications

Cross-participant within-modal and cross-modal classification analyses were conducted to determine whether affective properties of the video stimuli could be significantly predicted from fNIRS data within and across the watching and recall sessions.

In the within-modal classifications (Fig. 2), no significant decoding was observed for valence (M = .513, p=.224), arousal (M = .518, p=.148), or discrete emotion category (M = .26, p=.238) in the watching session. In the recall session, only arousal classification was significant (M = .537, p=.02), while classification of valence (M = .521, p=.117) and emotion category (M = .266, p=.149) did not reach significance.

**Fig. 2.**
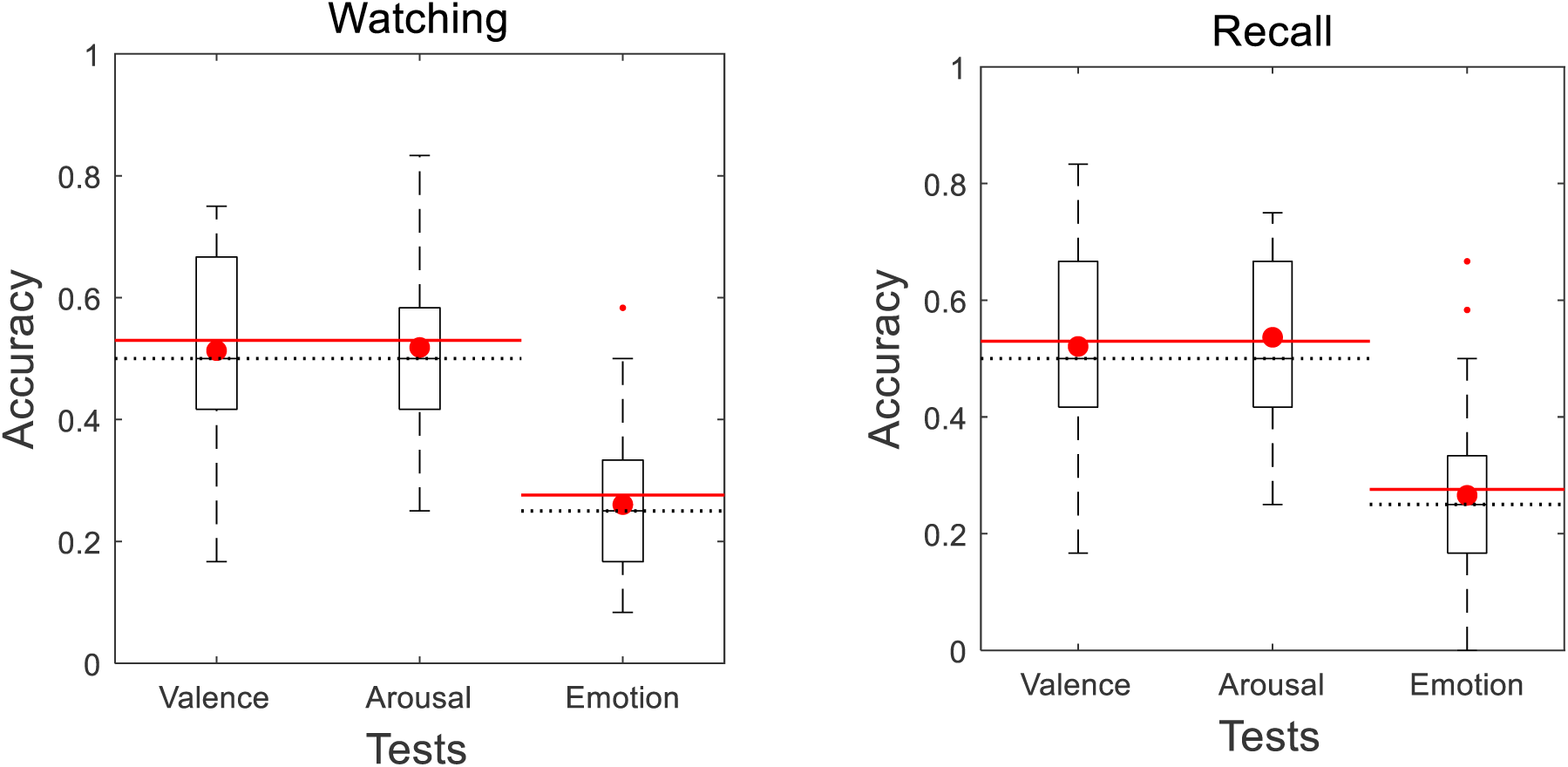
The boxplots of the cross-participant within-modal classifications within watching (left) and recall (right) sessions. The red dots represent the average classification accuracy values in the corresponding boxplots. The dotted lines indicate the chance level, and the solid red lines indicate the critical value.

**Fig. 3.**
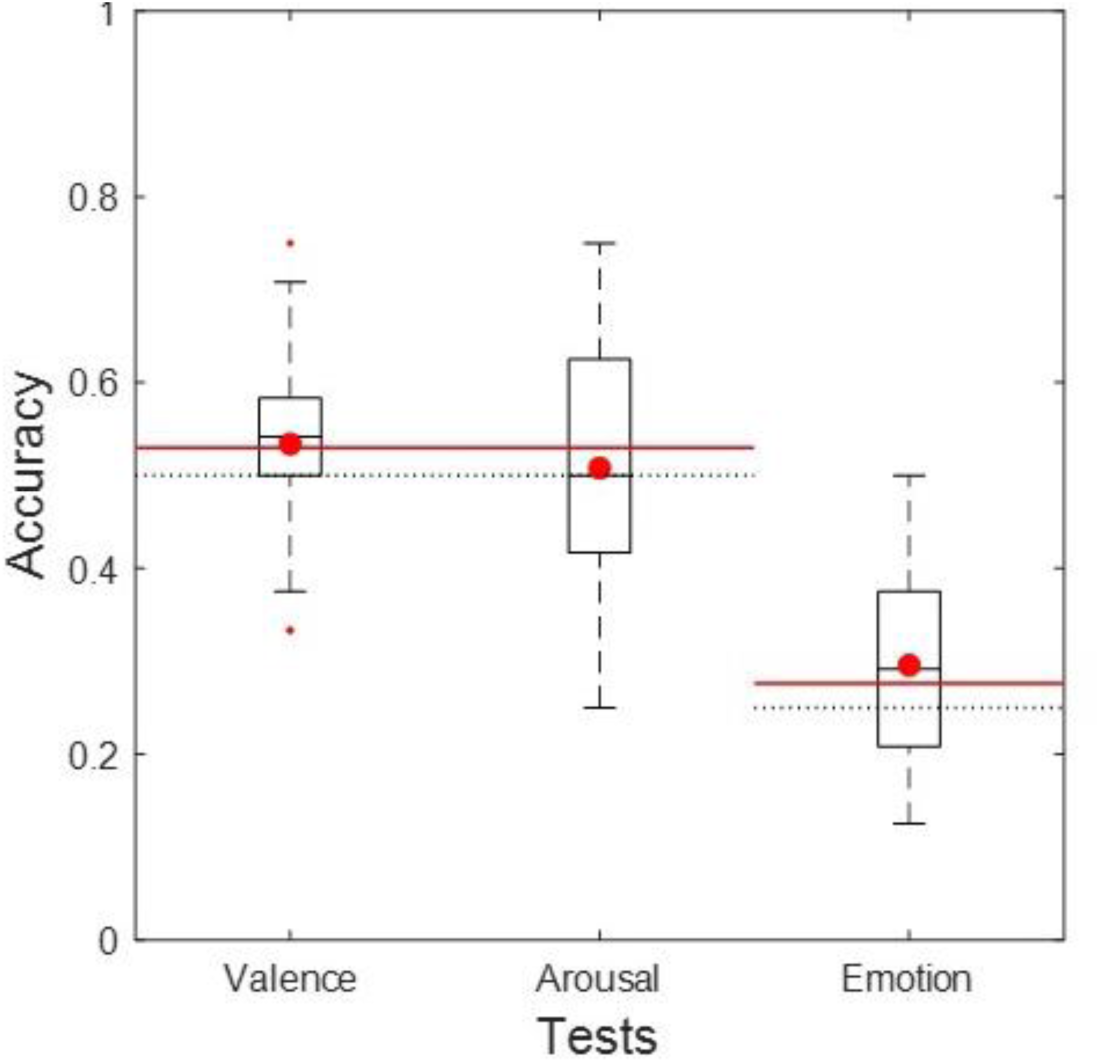
The boxplots of the cross-participant cross-modal classifications between watching and recall sessions. The red dots represent the average classification accuracy values in the corresponding boxplots. The dotted lines indicate the chance level, and the solid red lines indicate the critical value.

In the cross-modal classification, where training and testing were performed across different modalities (i.e., watching vs. recall), significant decoding was observed for valence (M=.534, p=.028) and emotion categories (M=.296, p=.002), but not significant for arousal (M=.501, p=.457).

Although decoding within each modality was limited, the significant cross-modal classification of valence and emotion categories suggests that these affective representations are preserved across perceptual and mnemonic states, supporting the presence of modality-general encoding.

## 4. Discussion

This study aimed to investigate how emotional experiences are represented in the brain during both the perception and recall of affective video stimuli. By employing fNIRS in a naturalistic setting, we examined whether core affective dimensions of valence and arousal could be decoded from prefrontal activation patterns across two distinct tasks including watching and recalling emotional stimuli. Using MDS and decoding techniques, we assessed the degree to which affective representations were stable across individuals and between perceptual and mnemonic states. Our findings revealed both shared and distinct neural representations of affect across the two modalities, suggesting a nuanced architecture of emotional memory processing in the prefrontal cortex.

Notably, our MDS results revealed significant effects for both modality-general and modality-specific dimensions of valence, but not for arousal. This result may seem contradictory, as modality-general significance suggests that emotional valence and arousal are encoded similarly during viewing and recall (Kim et al., 2017), whereas modality-specific significance implies that these representations differ across the two conditions. However, prior work suggests these patterns can coexist within a hierarchical coding scheme. For example, Shinkareva et al. (2014) demonstrated that while a supra-modal network in ventromedial prefrontal cortex encodes valence consistently across visual and auditory stimuli, each sensory modality also engages unique circuits for emotion processing. Similarly, Chen et al. (2017) found that high-level associative areas exhibit shared scene-specific patterns across perception and recall, yet also undergo systematic transformations during memory reconstruction.

One parsimonious interpretation is a two-stage hierarchical model of emotional memory. An initial, modality-general core affect signal preserves the broad valence of the original experience, followed by a secondary, modality-specific re-encoding during recall that adapts the memory to the demands of internal reconstruction (Skerry & Saxe, 2015). In this framework, the modality-general dimensions capture the invariant emotional gist, while the modality-specific dimensions reflect the dynamic, context-dependent elaboration that occurs when a memory is retrieved. Thus, the simultaneous significance of both contrast types does not indicate a true contradiction but rather reveals the brain’s capacity to maintain stable affective representations alongside flexible, task-dependent refinements.

Although one might expect that within-modal decoding (i.e., training and testing on the same session) would outperform or at least match cross-modal decoding (i.e., training on watching and testing on recall, or vice versa), our results showed the opposite. Within-modal classification was generally non-significant, whereas cross-modal classification achieved above-chance accuracies. At first glance, this pattern seems counterintuitive. In standard machine-learning terms, one would anticipate that models trained and tested on the same data distribution (within-modal) should yield higher accuracy than models tested on a different distribution (cross-modal) (Pereira et al., 2009).

However, two converging factors may explain our observation. First, the nature of emotional representation may be inherently more stable across the perceptual–mnemonic boundary than within each isolated context. Successful cross-modal decoding suggests that affective neural codes capture an invariant core that generalizes across the shift from viewing to recall (Kaplan et al., 2015). In contrast, within-modal decoding may suffer from session-specific noise, such as speech-related artifacts during recall or visual processing confounds during watching, that obscure identification of affective states when confined to a single modality (Pinti et al., 2018). Cross-modal classification, by conditioning on one modality and testing on the other, may effectively cancel out modality-specific noise, isolating the shared affective component (Nastase et al., 2016). Second, statistical power likely differed between within-modal and cross-modal tests due to the number of trials used in each. Within-modal classification for each session employed 12 trials per fold (one trial per emotion per participant), whereas cross-modal classification effectively doubled the data using 12 watching trials for training and 12 recall trials for testing, thereby leveraging 24 samples per fold. Cross-validation schemes are known to gain power as sample size increases, because the standard error of classification accuracy shrinks with more test cases (Hastie et al., 2009). Indeed, binomial tests for classifier significance become more sensitive when the number of observations grows (Combrisson & Jerbi, 2015). Thus, cross-modal decoding had a statistical advantage that may have elevated its significance relative to the underpowered within-modal classifications. Together, these considerations suggest that cross-modal decoding may outperform within-modal decoding when the increased number of trials enhances statistical power and the classifier is able to focus on a supra-modal affective template by filtering out modality-specific noise. This outcome aligns with recent fMRI work showing that cross-session or cross-task classification often reveals more abstract, stable representations than within-session decoding (Kaplan et al., 2015).

Our MDS results revealed both modality-general and modality-specific valence structures, whereas arousal lacked significant structural organization. Consistent with these findings, classification succeeded only for modality-general valence, while arousal decoding was restricted to modality-specific patterns, failing to generalize across conditions. This suggests that valence relies on a shared neural architecture across perceptual and mnemonic contexts, whereas arousal is encoded in distinct, context-sensitive representations. One possibility is that MDS and classification probe different aspects of the neural code. MDS captures graded similarity structure across all stimuli, revealing both shared and unique dimensions (Kriegeskorte & Kievit, 2013). The higher Pearson r for modality-general dimensions indicates that these contrasts explain variance in the similarity matrix, but they do not guarantee separability of individual trials by a linear classifier. Classification, by contrast, assesses the discriminability of trial-level patterns under a particular decision boundary (Mur et al., 2009). Thus, modality-specific variance may contribute to the overall similarity geometry without forming linearly separable clusters necessary for accurate classification within a single modality. Moreover, the relative magnitude of signal and noise differs between cross-modal and within-modal contexts. Within each session, modality-specific reconstruction processes during recall introduce idiosyncratic noise such as linguistic, motor, and imagery components that reduce within-modal classification accuracy (Penny et al., 2011). MDS, being unsupervised, integrates this noise into the overall distance structure but does not require clear separation for statistical significance testing. Cross-modal classification, by training on one modality and testing on another, may filter out these session-specific idiosyncrasies, allowing the core, modality-general affective signal to dominate (Nastase et al., 2016; Chen et al., 2017). In sum, MDS highlights the multifaceted representational geometry including both shared and unique dimensions, whereas classification reveals whether these dimensions support reliable trial-by-trial decoding within or across tasks. The combination of both methods indicates that emotional memory comprises a stable, cross-modal core embedded within a richer, context-dependent representational landscape revealed by MDS.

Our use of the OBELAB NIRSIT LITE system allowed us to sample four prefrontal subregions including IFG, MFG, dlPFC and mPFC. The IFG is known to support explicit emotion labeling and inhibitory control (Aron et al., 2004). Gao and Shinkareva (2021) identified the IFG as encoding both modality-general and modality-specific valence, which aligns with our findings: the IFG contributed to shared valence representations (modality-general) and also captured differences between perception and recall (modality-specific). The MFG is associated with working memory and attentional switching (D’Esposito et al., 1995), and has been shown to track arousal levels across contexts in affective tasks (Chikazoe et al., 2014). Similarly, the dlPFC has been shown to encode both modality-general and modality-specific valence (Gao & Shinkareva, 2021), consistent with our MDS results revealing both general and specific valence dimensions. Lastly, the mPFC plays a central role in autobiographical emotional recall (Buckner & Carroll, 2007).

Although Kim and Kim (2025) similarly investigated shared and unique representation of affective states between encoding and recall conditions, their actual fMRI results highlighted significant clusters in right middle temporal gyrus, right inferior temporal gyrus, and left fusiform gyrus, rather than prefrontal cortex. This anatomical discrepancy underscores both convergent and divergent aspects of emotion representation across imaging modalities. Convergent evidence arises in the shared affective dimension. Both our fNIRS study and Kim and Kim’s fMRI study (2025) found robust representation of valence across perception and recall phases, albeit in different cortical nodes. They demonstrated that temporal and fusiform circuits carry valence information consistently during encoding and recall, aligning with our cross-modal decoding of modality-general valence in prefrontal regions. This convergence supports models of distributed affect networks in which core valence signals propagate through both ventral temporal and prefrontal hubs (Lindquist & Barrett, 2012).

Divergent aspects reflect modality- and method-specific sensitivities. fMRI’s whole-brain coverage and high spatial resolution favored detection in ventral temporal areas, specialized for visual and semantic processing of emotional content (Peelen et al., 2010), whereas our fNIRS montage targeted prefrontal cortex, revealing how frontal control and self-referential networks maintain affective templates across tasks (Buckner & Carroll, 2007). The absence of prefrontal clusters in Kim and Kim (2025) may reflect fMRI’s relative insensitivity to superficial cortical dynamics during overt speech in recall, which fNIRS may capture in ecologically valid settings (Quaresima & Ferrari, 2019). In sum, Kim and Kim’s (2025) temporal and fusiform valence codes and our prefrontal valence and arousal codes may represent complementary nodes in an emotion-memory network. Future multimodal imaging combining fNIRS and fMRI could directly link these complementary representations.

Although our fNIRS study provides novel insight into modality-general and modality-specific affective coding in the prefrontal cortex, several methodological limitations warrant mention and suggest avenues for future research. First, fNIRS measures only cortical surface hemodynamics and cannot capture activity in deeper structures (e.g., amygdala, hippocampus) that are critical for emotion and memory (Scholkmann et al., 2014). Also, the spatial resolution of fNIRS and the limited number of channels (15 in the NIRSIT LITE) constrain our ability to localize fine-grained patterns within prefrontal subregions (Pinti et al., 2018). Superficial physiological signals such as scalp blood flow and systemic fluctuations may contaminate cortical measurements. Looking forward, future studies should integrate fNIRS with complementary modalities such as fMRI or EEG to achieve both depth and temporal precision, enabling direct linkage between surface and subcortical affective circuits (Cui et al., 2011; Chandrasekaran et al., 2021). Finally, longitudinal and clinical studies will determine how modality-general and modality-specific affective representations evolve over time and differ in populations with mood and memory disorders (Chi et al., 2020). By addressing these limitations, future work can build a more comprehensive, multi-level model of emotional memory that bridges perception, recall, and interpersonal sharing.

## References

Aron, A. R., Robbins, T. W., & Poldrack, R. A. (2004). Inhibition and the right inferior frontal cortex. Trends in Cognitive Sciences, 8(4), 170–177. 10.1016/j.tics.2004.02.010.

Badre, D., & D’Esposito, M. (2007). Functional magnetic resonance imaging evidence for a hierarchical organization of the prefrontal cortex. Journal of Cognitive Neuroscience, 19(12), 2082–2099. 10.1162/jocn.2007.19.12.2082.

Boas, D. A., Elwell, C. E., Ferrari, M., & Taga, G. (2001). Twenty years of functional near-infrared spectroscopy: introduction for the special issue. NeuroImage, 85, 1–5. 10.1016/j.neuroimage.2013.05.059.

Berboth, S., & Morawetz, C. (2021). Amygdala-prefrontal connectivity during emotion regulation: A meta-analysis of psychophysiological interactions. Neuropsychologia, 153, 107767.

Buckner, R. L., & Carroll, D. C. (2007). Self-projection and the brain. Trends in Cognitive Sciences, 11(2), 49–57. 10.1016/j.tics.2006.11.004.

Baker, J. M., Rojas-Valverde, D., Gutiérrez, R., Winkler, M., Fuhrimann, S., Eskenazi, B.,…& Mora, A. M. (2017). Portable functional neuroimaging as an environmental epidemiology tool: a how-to guide for the use of fNIRS in field studies. Environmental health perspectives, 125(9), 094502.

Chen, J., Leong, Y. C., Honey, C. J., Yong, C. H., Norman, K. A., & Hasson, U. (2017). Shared memories reveal shared structure in neural activity across individuals. Nature Neuroscience, 20(1), 115–125. 10.1038/nn.4450.

Chikazoe, J., Lee, D. H., Kriegeskorte, N., & Anderson, A. K. (2014). Population coding of affect across stimuli, modalities and individuals. Nature Neuroscience, 17(8), 1114–1122. 10.1038/nn.3749.

Chakravarthula, L. N., & Padmala, S. (2023). Negative emotion reduces the discriminability of reward outcomes in the ventromedial prefrontal cortex. Social Cognitive and Affective Neuroscience, 18(1), nsad067.

Combrisson, E., & Jerbi, K. (2015). Exceeding chance level by chance: The caveat of theoretical chance levels in brain signal classification and statistical assessment of decoding accuracy. Journal of Neuroscience Methods, 250, 126–136. 10.1016/j.jneumeth.2015.01.010.

Cui, X., Bray, S., Bryant, D. M., Glover, G. H., & Reiss, A. L. (2011). A quantitative comparison of NIRS and fMRI across multiple cognitive tasks. NeuroImage, 54(4), 2808–2821. 10.1016/j.neuroimage.2010.10.069.

Cui, X., Bryant, D. M., & Reiss, A. L. (2012). NIRS-based hyperscanning reveals increased interpersonal coherence in superior frontal cortex during cooperation. NeuroImage, 59(3), 2430–2437. 10.1016/j.neuroimage.2011.09.003.

D’Esposito, M., Detre, J. A., Alsop, D. C., Shin, R. K., Atlas, S., & Grossman, M. (1995). The neural basis of the central executive system of working memory. Nature, 378(6554), 279–281. 10.1038/378279a0.

Denny, B. T., Jungles, M. L., Goodson, P. N., Dicker, E. E., Chavez, J., Jones, J. S., & Lopez, R. B. (2023). Unpacking reappraisal: a systematic review of fMRI studies of distancing and reinterpretation. Social cognitive and affective neuroscience, 18(1), nsad050.

Fishburn, F. A., Ludlum, R. S., Vaidya, C. J., & Medvedev, A. V. (2019). Temporal Derivative Distribution Repair (TDDR): A motion correction method for fNIRS. NeuroImage, 184, 171–179. 10.1016/j.neuroimage.2018.09.065.

Gagnon, L., Yücel, M. A., Boas, D. A., & Cooper, R. J. (2011). Improved recovery of the hemodynamic response in diffuse optical imaging using short optode separations and state-space modeling. NeuroImage, 56(3), 1362–1371. 10.1016/j.neuroimage.2010.11.074.

Gao, C., & Shinkareva, S. V. (2021). Modality-general and modality-specific audiovisual valence processing. Cortex, 138, 127–137. 10.1016/j.cortex.2020.12.009.

Gower, J. C. (1975). Generalized Procrustes analysis. Psychometrika, 40(1), 33–51. 10.1007/BF02291478.

Harmon-Jones, E., & Gable, P. A. (2018). On the role of asymmetric frontal cortical activity in approach and withdrawal motivation: An updated review of the evidence. Psychophysiology, 55(1), e12879.

Halbwachs, M. (1980). The Collective Memory. Harper & Row.

Hastie, T., Tibshirani, R., & Friedman, J. (2009). The Elements of Statistical Learning (2nd ed.). Springer.

Kaplan, J. T., Man, K., & Greening, S. G. (2015). Decoding shared semantic space from neural patterns of movie viewing and recall. NeuroImage, 109, 317–326. 10.1016/j.neuroimage.2015.01.035.

Kensinger, E. A., & Schacter, D. L. (2008). Memory and emotion. In M. Lewis, J. Haviland-Jones, & L. Feldman-Barrett (Eds.), Handbook of Emotion (3rd ed., pp. 601–617). Guilford Press.

Kim, H., & Kim, J. (2025). Consistent neural representation of valence in encoding and recall. Brain and Cognition, 186, 106296. 10.1016/j.bandc.2024.106296.

Kim, H., Kim, J., Nashiro, K., & Mather, M. (2017). Neural correspondence between participants during movie viewing predicts episodic memory performance. Neurobiology of Learning and Memory, 138, 9–19. 10.1016/j.nlm.2016.12.002.

Kim, J., Shinkareva, S. V., & Wedell, D. H. (2017). Representations of modality-general valence for videos and music derived from fMRI data. NeuroImage, 148, 42–54.

Kiyokawa, H., & Hayashi, R. (2024). Commonalities and variations in emotion representation across modalities and brain regions. Scientific Reports, 14(1), 20992.

Kriegeskorte, N., & Kievit, R. A. (2013). Representational geometry: integrating cognition, computation, and the brain. Trends in Cognitive Sciences, 17(8), 401–412. 10.1016/j.tics.2013.06.007.

Lindquist, K. A., & Barrett, L. F. (2012). A functional architecture of the human brain: Emergent opportunities from the science of emotion. Trends in Cognitive Sciences, 16(11), 533–540. 10.1016/j.tics.2012.09.003.

Lindquist, K. A., Wager, T. D., Kober, H., Bliss-Moreau, E., & Barrett, L. F. (2012). The brain basis of emotion: a meta-analytic review. Behavioral and brain sciences, 35(3), 121–143.

Miskovic, V., & Anderson, A. K. (2018). Modality general and modality specific coding of hedonic valence. Current Opinion in Behavioral Sciences, 19, 91–97. 10.1016/j.cobeha.2017.12.005.

Mo, L., Li, S., Cheng, S., Li, Y., Xu, F., & Zhang, D. (2023). Emotion regulation of social pain: double dissociation of lateral prefrontal cortices supporting reappraisal and distraction. Social cognitive and affective neuroscience, 18(1), nsad043.

Nastase, S. A., Halchenko, Y. O., & Hasson, U. (2016). Cross-modal searchlight classification: methodological challenges and recommended solutions. NeuroImage, 124, 115–126. 10.1016/j.neuroimage.2015.09.037.

Ochsner, K. N., Bunge, S. A., Gross, J. J., & Gabrieli, J. D. (2004). Rethinking feelings: an fMRI study of the cognitive regulation of emotion. Journal of Cognitive Neuroscience, 14(8), 1215–1229. 10.1162/089892902760807212.

Pascalis, O., de Haan, M., Nelson, C. A., & de Schonen, S. (2013). Recognition memory for complex objects and faces in 3- and 6-month-old infants and adults: a neural network approach. Developmental Science, 16(1), 59–68. 10.1111/desc.12003.

Peelen, M. V., Atkinson, A. P., & Vuilleumier, P. (2010). Supramodal representations of perceived emotions in the human brain. Journal of Neuroscience, 30(30), 10127–10134. 10.1523/JNEUROSCI.2161-10.2010.

Penny, W. D., Friston, K. J., Ashburner, J. T., Kiebel, S. J., & Nichols, T. E. (2011). Statistical Parametric Mapping: The Analysis of Functional Brain Images. Elsevier.

Pereira, F., Mitchell, T., & Botvinick, M. (2009). Machine learning classifiers and fMRI: A tutorial overview. NeuroImage, 45(1 Suppl), S199–S209. 10.1016/j.neuroimage.2008.11.007.

Pinti, P., Tachtsidis, I., Hamilton, A., Hirsch, J., Aichelburg, C., Gilbert, S., & Burgess, P. W. (2018). The present and future use of functional near-infrared spectroscopy (fNIRS) for cognitive neuroscience. Annals of the New York Academy of Sciences, 1464(1), 5–29. 10.1111/nyas.13948.

Quaresima, V., & Ferrari, M. (2019). Functional near-infrared spectroscopy (fNIRS) for assessing cerebral cortex function during human behavior in natural/social situations: A concise review. Organizational Research Methods, 22(1), 46–68. 10.1177/1094428116658959.

Russell, J. A. (2003). Core affect and the psychological construction of emotion. Psychological Review, 110(1), 145–172. 10.1037/0033-295X.110.1.145.

Rolls, E. T., Huang, C. C., Lin, C. P., Feng, J., & Joliot, M. (2020). Automated anatomical labelling atlas 3. Neuroimage, 206, 116189.

Scholkmann, F., Kleiser, S., Metz, A. J., et al. (2014). A review on continuous wave functional near-infrared spectroscopy and imaging instrumentation and methodology. NeuroImage, 85, 6–27. 10.1016/j.neuroimage.2013.05.004.

Shinkareva, S. V., Wang, J., Kim, J., Facciani, M. J., Baucom, L. B., & Wedell, D. H. (2014). Representations of modality-specific affective processing for visual and auditory stimuli derived from functional magnetic resonance imaging data. Human brain mapping, 35(7), 3558–3568.

Skerry, A. E., & Saxe, R. (2015). Neural representations of emotion are organized around abstract event features. Current Biology, 25(15), 1945–1954. 10.1016/j.cub.2015.06.009.

